# A customizable tablet app for hand movement research outside the lab

**DOI:** 10.1101/633305

**Authors:** Adam Matic, Alex Gomez-Marin

## Abstract

**Background:** Precise behavioral measurements allow the discovery of movement constraints that provide insights into sensory-motor processes and their underlying neural mechanisms. For instance, when humans draw an ellipse on a piece of paper, the instantaneous speed of the pen co-varies tightly with the local curvature of the path. Known as the speed-curvature power law, this phenomenon relates to fundamental questions of motor control.

**New Method:** We have developed a software app for displaying static curves or dynamic targets while recording finger or stylus movements on Android touch-screen tablets. Designed for human hand movement research, the app is free, ready-to-use, open-source and customizable.

**Results:** We provide a template experimental protocol, and detailed explanations to use it and flexibly modify the code for different kinds of tasks. Our validation of the app demonstrates laboratory-quality results outside the laboratory. We also provide raw data and analysis scripts.

**Comparison with Existing Methods:** Commonly used laboratory devices for recording hand movement trajectories are large, heavy and expensive. In turn, software apps are often not published, nor customizable. Our app running on tablets becomes an affordable, flexible, and portable tool suited for quantitative and robust behavioral studies with large number of participants and outside the laboratory (e.g. in a classroom, a hospital, or at home).

**Conclusions:** The affordability, flexibility, and resolution of our tablet app provide an effective tool to study behavior quantitatively in the real world.

**Highlights:**

- A free, ready-to-use, open source, and customizable app for Android tablets.
- High-resolution measuring of finger movement during tracing, tracking & scribbling.
- Fast and easy data collection and experimental design with affordable hardware.
- Allowing for high-throughput experiments outside the lab (classroom, hospital, home).
- Validated for state-of-the-art research (speed-curvature power law, drawing accuracy).

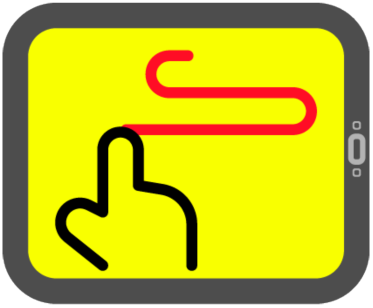

## 1. INTRODUCTION

It has been argued that nothing makes sense in neuroscience except in the light of behavior (**Krakauer et al, 2017**). Yet, even when we carefully measure the behavior of organisms, the promise that the discoveries found in the laboratory will generalize in real-world situations is often hard to fulfill. This is in part due to the simplicity of experimental designs which, in turn, allow to maximize control by the experimenter, taming the complexity and context that is natural to the behaving subject under study (**Gomez-Marin & Mainen, 2016**). For instance, writing on a piece of paper or simply drawing with our finger on a tablet are everyday activities, but the quantitative study of the processes and mechanisms generating such complex hand trajectories is nearly always done in laboratory conditions.

Another main reason for laboratory research is the necessary involvement of expensive, sophisticated, and usually massive technological devices for manipulation and measurement. In fact, measuring behavior has a rich history in hand movement research, where the development of recording instruments has played a central role. These include graph paper, cameras, robot arms, motorized linkages, and other clever gadgets to store hand position over time. While it is not our aim here to present a exhaustive account, let us list several influential methods in movement research that illustrate the advancement of a field with more than a century of history.

An early instrument in recording movement was the Edison pen, where a needle at the tip of the pen was oscillating at constant frequency. The needle made marks on the paper so that movements at higher speed left marks spaced further apart than movements at lower speed. With such device, the speed of the pen in curved parts of a trajectory was observed to be lower than the speed in straight parts (**Binet & Courtier, 1893**). A middle-sized model was priced at 50$ in the 1890’s, which is on the order of 1500$ in today’s dollars. A few years later, Woodworth used a simple method of graph paper and metronome-synchronized movements to measure the relationship between speed and accuracy (**Woodworth, 1899**). As he notes, the method was easy to use and he recorded more than 125K individual trajectories for a study. The difficult part was analyzing the recorded data, and this was done by his assistants. During the 1930’s, Bernstein invented a highly sophisticated method called cyclography, utilizing high-speed film cameras with shutter speeds of 150-200Hz and light-bulb markers placed on the bodies of his participants (**Bernstein, 1984; Gurfinkel & Cordo, 1998**). Using multiple cameras or a single camera and a system of mirrors, the three-dimensional trajectories of joints and limbs of participants could be reconstructed. Bernstein formulated the so-called degrees of freedom problem, and an early theory of movement control hierarchy. Regarding handwriting analysis with digitizers in the 1960’s, an overview of devices used can be found in (**Schoemaker, 1998**).

In the 1980’s, a puzzling constraint between instantaneous speed and local curvature of end-point hand trajectories was discovered in data recorded with an ultrasonic device called the Graph Pen (**Lacquaniti, Terzuolo & Viviani, 1983; Soechting, Lacquaniti & Terzuolo, 1986**), which was capable of 100Hz sampling and 0.1mm accuracy in measuring pen position on a plane. Another device used in (**Lacquaniti, Terzuolo & Viviani, 1983**) was a Calcomp electromagnetic digitizing table, 100Hz sampling and 0.025mm accuracy. It was then established that in human hand movement, the instantaneous angular velocity is proportional to the local curvature raised to the 2/3 power (A=k · C^β^ with β=0.66); the so-called two-thirds power law.

Further investigations of the coordination of arm movements used two-link mechanical manipulanda. Built with two joints and precision potentiometers calibrated to measure joint angles, and sampling at 100Hz, it achieved 1mm resolution in the endpoint position measurement (**Flash & Hogan, 1985**). Another class of measuring technologies consisted of pressure sensitive pads, which can be used with ordinary pens. For example, a Quest Micropad pressure sensitive device can reach 200Hz, and achieve 0.2mm accuracy (**Wann, Nimmo-Smith & Wing, 1988**). Furthermore, for free movement in three dimensions, the use high-speed cameras together with visual markers placed on the body of the participant facilitate computerized analysis. For instance, in (**Dounskaia et al., 2002**) an Optotrack 3D optoelectronic camera system achieved 100Hz frequency using infrared LED lights as markers.

More recently, researchers have been using digitizing graphics tablets like the Wacom Cintiq and Intuos. In particular, using such devices it has been empirically found (and theoretically predicted) that humans produce a spectrum of speed-curvature power laws while tracing pure frequency curves (**Huh & Sejnowski, 2015**). These devices provide very high temporal and spatial resolution of recording pen or finger position, up to 140Hz in sampling rates for Cintiq and up to 200Hz for Intuos models, and reported 0.005mm of spatial resolution (5080 lines per inch; but accuracy may be lower), while displaying any curve geometry and target kinematics on the very surface where the participant draws. We have recently reproduced such findings with the same devices (**Zago, Matic et al., 2018**). However, note that the Wacom Cintiq 27QHD is a 27” monitor weighing 13kg, and priced around 2750$. Its size and cost, and the requirement of a separate computer to record the data can be a limitation in experimental settings that require affordable, portable, and high-throughput data collection. This has prompted us to explore other solutions that are more efficient and inexpensive without compromising the quality of the data.

In the last years, small-size autonomous computers such as iPad tablets, Android tablets, touch-screen laptops or even smart-phones are increasingly used in movement control and development research (**Accardo et al., 2013; Lee et al., 2014; Hill et al., 2014**), as well as in clinical settings (**Anzulewicz et al., 2016; Sisti et al., 2017; Vianello et al., 2017**). Tablet computers are affordable and transportable, which in principle makes them ideal for large-scale experiments outside of the laboratory, in natural settings for humans such as classrooms, homes, or hospitals. Actually, the spatial and temporal resolution of recording movement trajectories in the tablets is becoming on par with larger, specialized graphics digitizing tablets, thus becoming a reasonable and practical alternative. We have exploited this fact here.

In this article we report on creating a free and open source application for an Android tablet made to facilitate large-scale hand movement experiments in situations not necessarily constrained to laboratory settings. Our application can be used in its current form, or as a template and code base for designing applications for new experiments. Currently, there are three main task types available in the code: (i) tracing figure shapes, (ii) tracking target trajectories and (iii) free scribbling or drawing. Each task type invites to constraint certain aspects of trajectory production. For instance, in tracing, participants are invited to move their finger following a particular geometry statically displayed on the screen, but with kinematics being free. In tracking, participants are invited to follow the particular kinematics displayed by a moving target. And in scribbling, participants can draw in space (geometry) and time (kinematics) as they please. We have developed an experimental protocol for high-throughput experimentation outside the lab, and we have tested the validity of the app for generating laboratory-quality motor control data. In sum, our app is ready to use, open, customizable, and suitable for human movement research.

## 2. MATERIALS AND METHODS

### Hardware

We have used a fairly common and affordable tablet, the Samsung Galaxy Tab A6 (alternative name SM-T580) whose price is around 170€. Physical dimensions are 254 × 164 × 8mm. It comes with the Android operating system, version 8.1.0 (Oreo) and API level 26. The display is a 10.1” PLS LCD screen, with dimensions 216 × 135mm, and a resolution of 1920 × 1200px. The tablet has a capacitive touch-screen, and registers touch by a finger or a capacitive stylus, with resolution equal to the display resolution, which is 226ppi or 8.89 px/mm in pixel density. Maximum screen refresh rate is 60Hz. Maximum sampling rate of touch events is not published, but we have found it to be close to 85Hz.

### Software

The app was programmed in Android Studio (version 3.3.2), a free integrated development environment (IDE), officially supported by Google, intended for development of Android OS applications on multiple platforms. Android Studio enables development in programming languages Java, C++, Go and the language we used for this application was Kotlin. Kotlin is a recently designed general-purpose programming language fully interoperable with Java, can freely use Java libraries, and compiles to the JVM, but features a simpler and more concise syntax. The combination of relative simplicity and the ability to use existing Java libraries makes Kotlin a practical choice. To program the app, we have used a Windows 10 PC, with 8GB of RAM and Intel i5 CPU. But there are no stringent constraints on the PC needed to do so.

### Experiments

The proof-of-concept validation behavioral experiments were performed by one of the authors. They involved tracing, tracking and drawing different geometric and kinematic tasks with the finger on the tablet. The total duration of the experimental protocol coded in the app was 15 minutes. Default instructions were to produce fast and fluid movements without corrections. Procedures were approved by the Institutional Review Board.

## 3. RESULTS

We have created an application software (an “app”) for Android tablets to be used in hand movement and sensory-motor control research, with a focus on the speed-curvature power law. The application is ready for use in its current form. We also provide the source code together with an easy way of designing other tasks to be encoded in the app, as well as a deconstruction of an effective experimental protocol, which we demonstrate. We also validate the quality of the data collected for motor control science, and share the raw data as well as analysis scripts. See **Figure 1** for a general methodological scheme, whose steps we now explain:

**Figure 1.**
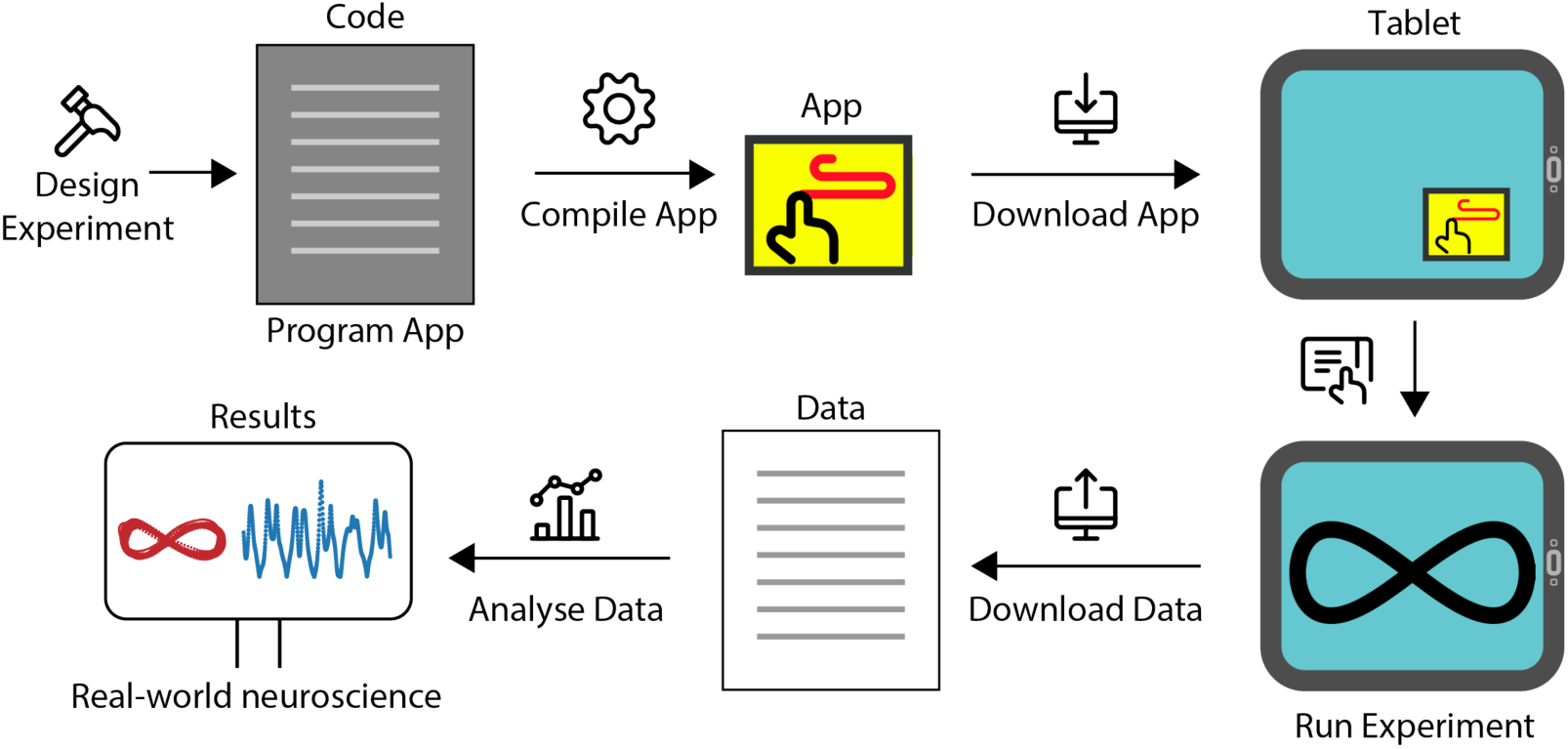
General methodological scheme of the tablet app: from experimental design to data analyses. The app is compiled and ready to be used (*app-release.apk* file). It simply needs to be downloaded to the Android tablet via USB from a desktop computer or by email. Its deployment consists of three phases: basic data entry, practice and experiment sections (see **Figure 2**). When the experiment is finished, the raw data (movement trajectories, experimental templates, and participant metadata) can again be easily transferred from each tablet to a desktop computer via USB or email. It is also possible to customize the app for other experimental designs involving sequences of tracing, drawing or scribbling tasks (see **Figure 3**). This is implemented in the source code of the app (editing the *Experiment.kt* file; see scripts in supplementary material). Quantitative data analyses (which can be performed in Python files we share within a Jupyter Notebook; see *Analysis_KinematicCognition.ipynb* file in supplementary material) yield state-of-the-art motor control results as demonstrated in **Figure 4**. The potential of the app for real-world behavioral neuroscience experiments is summarized in **Figure 5**.

### 3.1. The App is ready to run and easy to install

The application can be installed on any Android tablet. It can be run in its original form by simply downloading it in a tablet. This would install the app with predesigned template experiments and its default settings. An ordinary route for Android applications is the Google Play Store, but it is not necessary, as it might involve fees and delays, and add another layer of complexity to the process. Distributing the *apk* file can be done via USB cable, copying it from the PC to each tablet, or more simply via email, by sending the *apk* file or a link for its download to each tablet’s email address. The app can then be downloaded and installed on the tablets.

### 3.2. An effective experimental protocol has been designed and validated

As diagrammed in **Figure 2**, when participants start the app the first panel they see is the data entry panel. They are asked to enter the year and month of birth, gender, and dominant hand. This information is stored as metadata, and used to construct the filename with the trajectory data. The participant can then start a “practice” sequence or an experiment sequence, following the instructions previously programmed in the app by the experimenter. The practice sequence serves to familiarize the participant with the tasks and can be repeated as many times as needed. By default, the data is not recorded in the practice sequence (but this can be modified in the *Experiment.kt* file; see next subsection). The “experimental” sequence contains a series of tracing, tracking and scribbling tasks, as defined in the program. Data is recorded after each task in the experimental sequence. The experiment ends after all tasks have been run.

**Figure 2.**
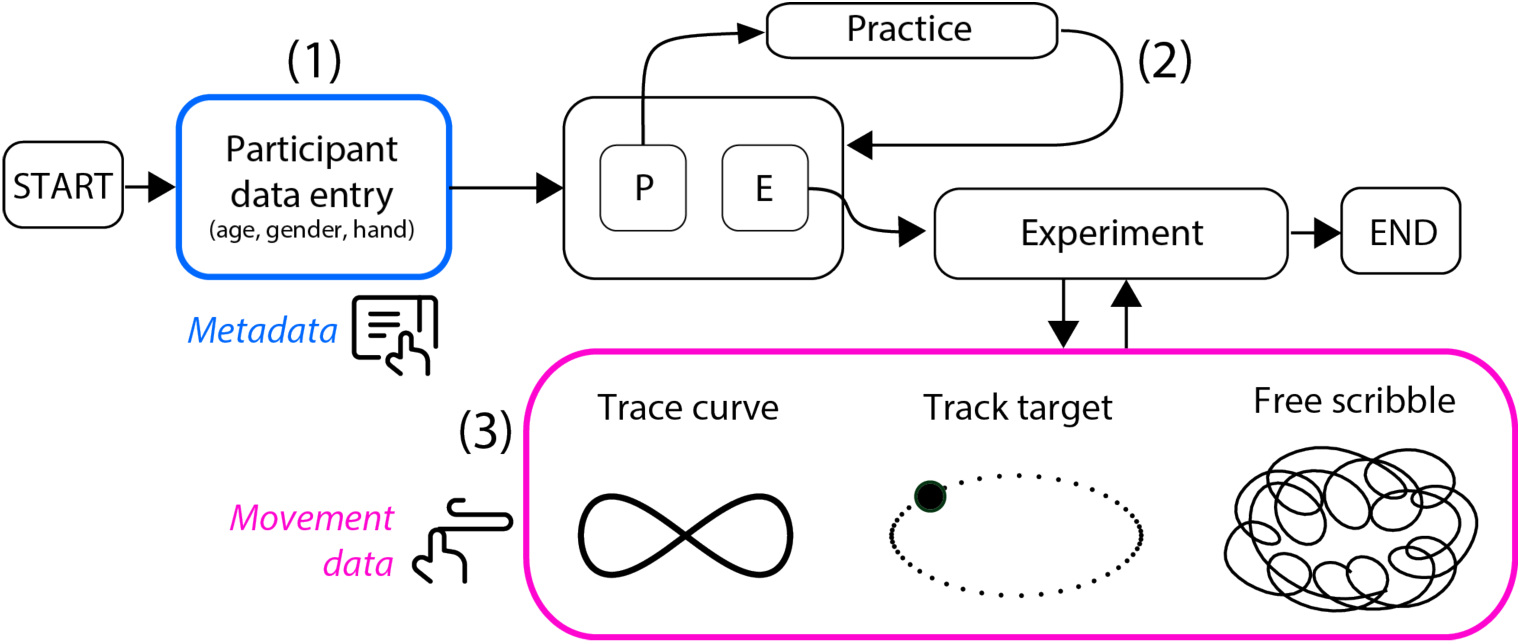
Application flow diagram during an experiment. Upon clicking on the app icon on the tablet desktop, the app starts with the data entry panel (**1**), which is saved as metadata. Next the participant can choose to run a practice sequence (**2**) one or more times by pressing the “practice” button (**P**). Then the participant may click the “experiment” button (**E**) so as to be presented with a sequence of tracing, tracking and scribbling tasks (**3**) for which data is recorded.

After a concise verbal instruction to the participant about the experiment, we found practice to be important in ensuring that the experiment takes place smoothly. We also found that it is effective to present the various (tracing, tracking, and scribbling) tasks consecutively with a brief pause, rather than providing a general menu where the participant clicks back and forth the corresponding task or curve to execute. This protocol coded in the app should be particularly useful to perform high-throughput experiments in groups of children or adults by having the app installed in several tablets and running the practice and experiment phases synchronized across participants.

When the experimental sequence is finished, the application ends and the movement data (x position, y position, time) is saved in a *txt* file together with the type of task and metadata (age, gender, hand) as the file name, so that each file self-contains all the necessary information for further analyses (see section 3.6).

### 3.3. The app can be edited by non-professionals to customize experimental protocols

The source code is provided and can be edited in order to accommodate the particular needs of the experimenter. To facilitate the customization process, we have designed the code to allow editing of a single file in order to change the most important protocol elements: the type of task, duration, sequence of appearance, and pause in between tasks.

After the experimental design is implemented in code, tested, and debugged, the code needs to be compiled into an Android package file (*apk*) using the Android Studio IDE. Note that the code is written in the Kotlin programming language. To make it accessible to nonprofessionals, the file *Experiment.kt* in the project source is the only one that needs to be edited (see supplementary material). It contains definitions of all the curves used in tracing tasks, all target trajectories used in tracking tasks, the duration of each task and their ordering in the practice sequence and the experiment sequence.

The definitions of the curves are at the top of the file *Experiment.kt*. Curves for tracing tasks, such as ellipses or lemniscates, are defined as lists of x-y points. For future reference and comparison with participant trajectories, points for each curve are saved into a text file named after the curve (e.g. *Lemniscate.txt* contains a list of x-y points used to draw the shape on the screen). Target trajectories are defined as functions that return point coordinates at a particular time t measured from the task start. Currently implemented code enables design of target trajectories following pure frequency curves (specifying geometry) and velocities defined by a speed-curvature power law with an arbitrary exponent (specifying kinematics).

Next, each task or event needs to be defined with a name, type and duration. The name is arbitrary, the type is one of “trace”, “track”, “scribble” or “pause”, and duration is the number of seconds after which the task will automatically end and proceed to the next task. Finally, ordering and duration of tasks are defined for the practice sequence and the experiment sequence.

In sum, in order to customize the app, one needs to download the project from Github (clone the repository), and open it in Android Studio to edit the file named *Experiment.kt*. As depicted in **Figure 3**, this allows a handy composition of new “practice” and “experiment” tasks.

**Figure 3.**
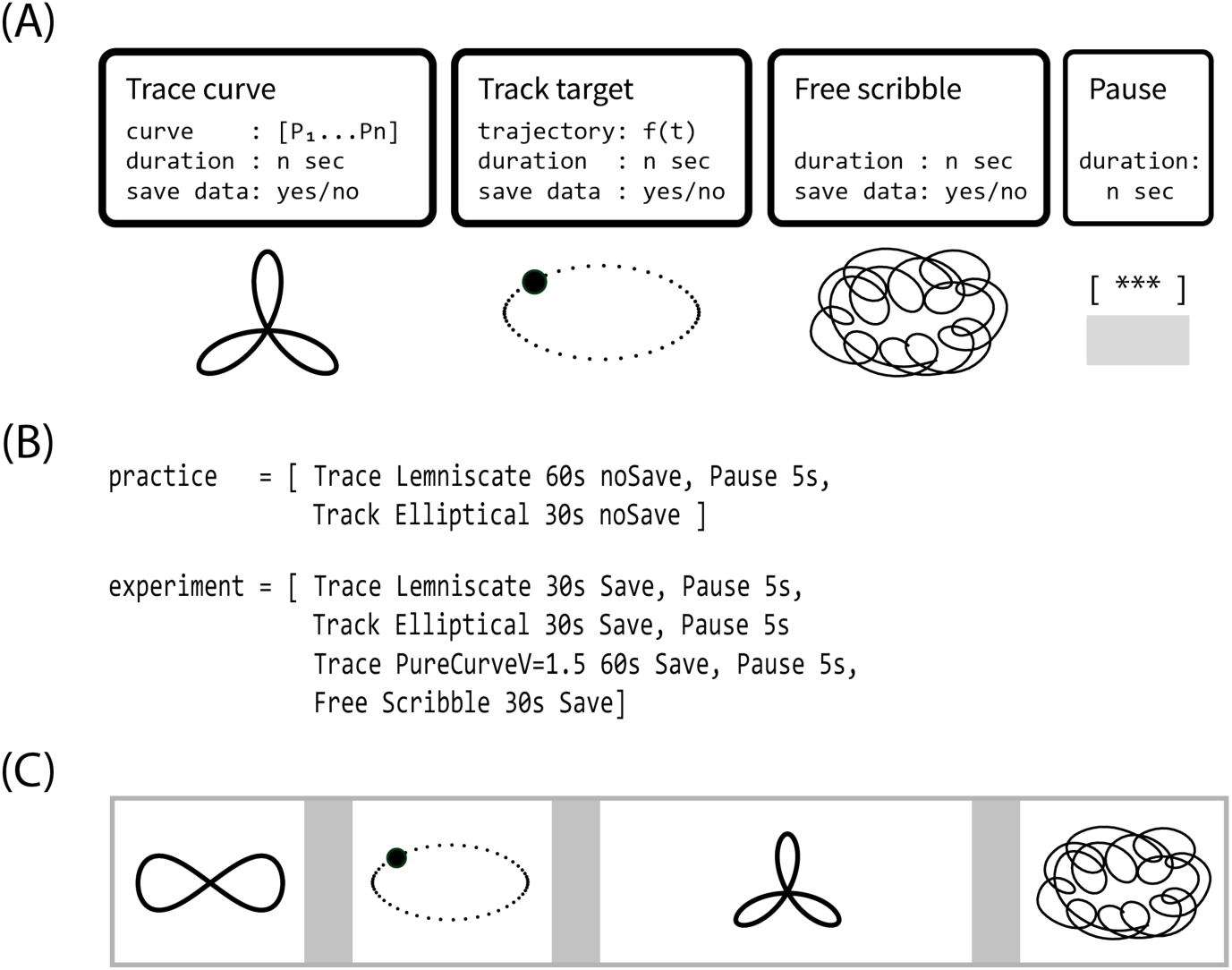
Composing task elements to easily and flexibly create an experimental protocol in the app. The *Experiment.kt* file contains: (**A**) the definitions of curves and trajectories to specify the geometry and kinematics used as experimental tasks, as well as definitions of each task specifying duration and whether to save the data or not; and (**B**) definitions of practice and experimental sequence of tasks as one wishes to make them appear in the application. For instance, the “experiment” vector in (B) would generate the sequence of tasks depicted in (**C**).

### 3.4. The app can be thoroughly customized by advanced programmers

We provide all the necessary source-code files as Supplementary Material. In particular, one needs to access the “KinematicCognition” folder. The files therein (and also inside the “idea” folder) are the build instructions for Android Studio and configurations for the project. They are mostly in Kotlin programming language. In the “gradle” folder one finds additional files for the building process. There is no need for the user to modify any of these files. The Android Studio actually generates and modifies them as one compiles the app. In the “app” folder one finds two main folders. In the “release” folder one can find the app ready to be installed as an *app-release.apk*. The “src/main” folder contains all the scripts needed to customize the app. In the “java/com/example/kinematiccognition” are the Kotlin (.*kt*) files corresponding to the so-called ‘activities’ (screens, routines for recording the trajectories, saving files, generating trajectories). For basic editing as described in the previous section, one does not need to worry about any of such files. But advanced programmers can of course make use of their editing. In the “res” folder there are many folders automatically managed by Android Studio. They comprise icons, layouts of the screens, connections between layouts, additional libraries, and dependencies. Let us also remind to select the appropriate API level for compilation in Android Studio so that it matches the particular tablet model to be used. Note that if the application is intended for a tablet with different screen resolution (ours was 1920 x 1200px), the shapes and trajectories should be adapted by adjusting their size in pixels in *Experiment.pk* file.

### 3.5. The app is optimized for temporal resolution of trajectory recording

In the Android operating system, the touch location and the timestamp are not usually provided in their raw form, as recorded by the touch-screen driver. To improve user experience during normal use, finger touch locations are by default recorded in batches of events, synchronized to display refresh events, and passed through an interpolation and estimation algorithm. These touch events are available to the programmer through methods *event.X, event.Y* for the location, and *event.getTime* for the timestamp. Maximal temporal resolution is equal to the screen refresh rate, which is 60Hz. These methods are useful in general user interface programming, gesture recognition and similar uses. However, the interpolation and estimation algorithms may distort finger touch position and timestamp. Similarly, because the touch events will be synchronized to screen refresh events, the rate of touch events may be lower than recorded in its raw form.

To acquire more accurate and non-processed raw location and timestamp data at maximal possible temporal resolution, we access the recorded batches of events through the *event.historicalX, event.historicalY* and *event.historicalTime* methods. Trajectory recording methods are implemented in each of the task classes in the code. In target tracking tasks, trajectories are defined as functions of time. This method allows for correct positioning of the target, independent of the drawing frame rate or lags in the running of the app during the task. Synchronization of the target and finger trajectories in data analysis can be made using this target trajectory data. In free scribbling tasks, only the last one second of the trajectory is shown, as a disappearing trail. This minimizes the effect of drawing on the frame rate, keeping it near maximum 60Hz.

### 3.6. Movement data and metadata file formats allow efficient management and analysis

Each tablet will contain the data of the experiments that were run on it, stored by default to the folder “/internal storage/download”. The data is composed of text files containing participant trajectories, target trajectories, and default curve points. As we mentioned, they can be copied to the desktop computer over a USB cable, or sent to an email address from each tablet.

The filename of each recorded trajectory contains the metadata of the participant information collected in the data entry panel (year and month of birth, gender, and dominant hand), as well as the type of the task performed, and the time and date of the experiment. For example, file *February1986MaleRight scribble 10.4.2019. 16.10.57.txt* contains the movement data of a scribbling task performed at the noted date and time by a right-handed male born in February 1986. In this way, all the relevant information of each experiment is centralized in a single file.

Raw trajectory data is stored in text files, with each file containing three columns, a timestamp in milliseconds since the start of the task, and x and y coordinates in pixels. Note that the upper left corner is the coordinate (0, 0), x is increasing from left to right, and is y increasing from top to bottom. This may result in reversing the y coordinate if the data is plotted in the traditional Cartesian coordinate system.

Curve tracing and scribbling tasks save the participant movement coordinates only, while the tracking tasks save two files: one with participant data, with the filename prefixed “user”, and one with target positions prefixed “target”. Target participant data are saved in different files because of their different sampling rate. Target position is saved at the rate of screen refreshing, while the participant data at the rate of touch event recording. For the tablet Samsung T580 used in developing this application, the timestamp differences were are approximately 16.66ms (60Hz refresh rate) for screen refresh, and 11.8ms (85Hz sampling rate) for touch events. While the rate of data sampling for participant trajectories is reasonably constant at near 85Hz, it is useful to spline/interpolate and re-sample the participant and target trajectory data, or participant data from different tasks to the same sampling frequency. For target tracking tasks, the target trajectory can be synchronized to participant finger trajectory by the timestamp variable, since the timestamps measure time in ms since the start of the task, for both movements.

### 3.7. The data collected with the app yields state-of-the-art scientific results

To evaluate the data collection potential of the app and to demonstrate the range and quality of possible types of analysis, we performed a pilot study consisting of several tracing, tracking and scribbling tasks. All data was filtered with a low-pass Butterworth filter with a cutoff frequency of 8Hz. The analyses we performed are characteristic of the study of the speed-curvature power law, as well as of other quantitative aspects of movement research. The results, shown in **Figure 4**, illustrate the usefulness of our method in hand movement research.

**Figure 4.**
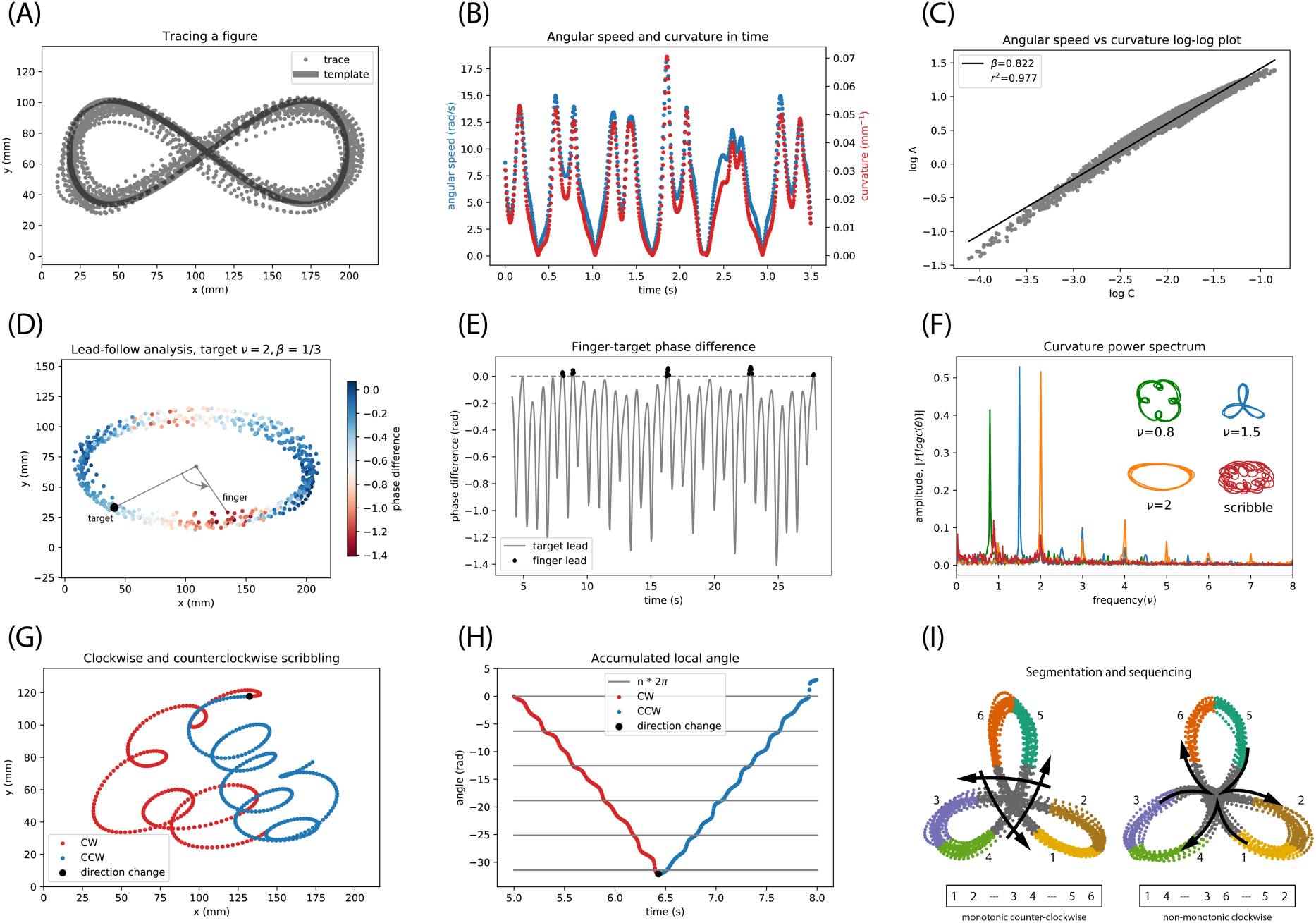
Analyses of the data collected with the app produce state-of-the-art results. *Five main situations are shown: power-law constraint during tracing (****A-C****); lead-follow dynamics during tracking (****D-E****); geometrical accuracy in pure-frequency curves (****F****); clockwise scribbling (****G-H****); and action segmentation degeneracy (****I***). **(A)** Tracing a lemniscate figure with the finger on the tablet. **(B)** Instantaneous angular speed and local curvature as a function of time for a short interval. Both appear tightly correlated. **(C)** The trajectory of the participant’s finger complies with the speed-curvature power law (r^2^ = 0.977), with an exponent β=0.82. **(D)** Tracking a moving target along an ellipse. The color of the dots depicts the relative phase angle (measured from the center of the ellipse) between target and the finger, which is minimal in the most curved parts of the trajectory. **(E)** Lead-follow analysis reveals that the participant is behind the target most of the time, only leading in front of the target at some points that coincide with maximal curvature. **(F)** Amplitude of the power spectrum of the curvature profile in tracing pure frequency curves shows strongest peaks at the frequency of the template, while the scribble has a much broader distribution. **(G)** Direction analysis during free scribbling shows clockwise turning (in red) for the first part of the analyzed trajectory, followed by counter-clockwise turning (in blue) in the second part. **(H)** Accumulated angle over time reveals five full windings before changing direction. **(I)** Discrete segmentation of a continuous path produced while tracing a three-lobe flower-like shape can reveal different choice sequences in drawing of the same path across different trials or individuals. Apart from clock-wise or counter-clockwise directions, one can also choose to trace the pattern with monotonic changes in curvature, or sharply changing direction at the center.

First, when tracing of a lemniscate figure (**Figure 4A**), the trajectory shows a strong covariance between angular speed and curvature (**Figure 4B**), which yields a power law with the exponent β=0.82 and r^2^=0.977 (**Figure 4C**). This is consistent with the law and exponent found in the literature for a lemniscate (**Viviani & McCullom, 1983**). Other curves tested (data not shown) yielded power laws with the exponents reported in (**Lacquaniti, Terzuolo & Viviani, 1983**) and (**Huh & Sejnowski, 2015**).

Second, we analyzed the lead-and-follow dynamics when the finger tracks a moving target along an elliptical trajectory with hypo-natural kinematics (**Figure 4D**). Hypo-natural movement trajectories are defined as those for which the angular speed and curvature power law has an exponent lower than 2/3 (in this case we imposed β=1/3) so that the target slows down in high-curvature parts of the path much more than in the movements naturally performed by participants. The angular difference between the target and participant positions is measured from the center of the ellipse at each point along the trajectory. Consistent with a similar analysis in the literature (**Viviani & Mounoud, 1990**), we find that the participant is not merely following the target, but getting closer and further away periodically, with more difficulty to track it at certain regions, and with certain trajectory segments even overtaking the target (**Figure 4E**).

Third, a set of pure frequency curves (**Huh, 2015**) with parameters ν=0.8, ν=1.5 and ν=2.0 (respectively corresponding to four-lobe, three-lobe and ellipse curves) were shown on the tablet screen as static templates and the participant traced those figures in a fast and fluid manner. For participant traces of those curves, **Figure 4F** shows the amplitude of the curvature spectrum, which is the Fourier transform of the logarithm of the curvature profile but parametrized in angle rather than in length or time (**Huh & Sejnowski, 2015**). Remarkably, the curvature profiles of the traced trajectories have single peaks at the precise pure frequencies of the template curves displayed. One also sees harmonics. In contrast, the log curvature profile of a free scribbling trajectory does not show sharp peaks (except some dominant contributions at ν=2 and also a bit below ν=1) as it is not a pure-frequency curve. Overall, this analysis illustrates how such spectra can be a powerful and principled measure of geometrical accuracy during tracing.

Fourth, in a segment of scribbling movements (**Figure 4G**) we examined the direction of movement as the accumulated unwrapped local angle over time. We can clearly distinguish between clock-wise and counter-clockwise movements, and quantify the number of complete rotations (gray lines in **Figure 4H**) during free scribbling.

Fifth, we can discretize a continuous trajectory by means of a segmentation analysis. As shown in **Figure 4I**, there are actually different ways to draw the same simple figure. The three-lobe pattern helps illustrate such degeneracy. In the left one, the trajectory crosses the center without changing the direction of movement (thus, monotonically) and this is all done counter-clockwise. In the right one, each ‘petal’ is drawn separately (non-monotonic curvature changes) with the direction of movement changing in the middle of the figure, while this is done clockwise. In sum, the tracing of such a simple figure can betray handedness and decision-making differences across participants, and within participants in time.

### 3.8. Scripts for data analyses are available as a Jupyter Notebook

The raw data and a python scripts to analyze it are also available as supplementary information. In particular, the “KinematicCognition-Analysis” folder contains the files *power_law_analysis.py* and *power_spectrum.py* which correspond, respectively, to the scripts that estimate speed and curvature to test the power-law constraint and it exponent, and the scripts that calculate the power spectrum of any trajectory. The file *Analysis_KinematicCognition.ipynb* is a Jupyter Notebook that facilitates the visualization and generation of the analyses corresponding to those shown in each plot of **Figure 4**. In the “data-new” folder is the raw data of the pilot study corresponding to different curves traced, targets tracked and scribbling. Note that for local use, one must match the paths to local folders.

## 4. DISCUSSION

Android tablets are widely available and affordable today. The reader may even have one or two at home. We have created an app and deployed it on a commercial tablet, demonstrating that it allows sufficiently high temporal and spatial resolution for state-of-the-art motor control laboratory research. We have provided a ready-to-use version of the app, with an experimental protocol design that is suited for high-throughput effective data collection outside the laboratory. We have also shared the source code and organized it so as to facilitate researchers to design their own experiments, and to be able to compile new app versions.

As summarized in **Figure 5**, our app has plenty of advantages. Let us also remark now some of its limitations. First, it is worth stating that the spatial and temporal resolution of digitizing tables and similar devices a few decades ago was on the same level (or better) than today’s Android tablets. For instance, in (**Lacquaniti et al., 1983**) a resolution of 100Hz is reported for an electromagnetic digitizing table with 0.025mm of accuracy, and in (**Wann et al., 1988**) a pressure sensitive pad achieved 200Hz sampling rate, though with somewhat lower accuracy of 0.2mm. Nevertheless, the sampling rate of movement recording for the tablet used in the present manuscript is fast enough to enable the quantitative analysis of kinematics and geometry of human movement while drawing. The price of the tablet we used here is easily an order of magnitude cheaper than typical recording devices used in the lab. Second, we did not program the app to be run on cell phones since this would considerably limit the spatial range and resolution of most of the motor control experiments one is interested in. Third, note as well that our app cannot be run on an Apple iPad. Yet, nothing prevents other users to use the code, design, and analyses employed here to extent it to other platforms or uses.

**Figure 5.**
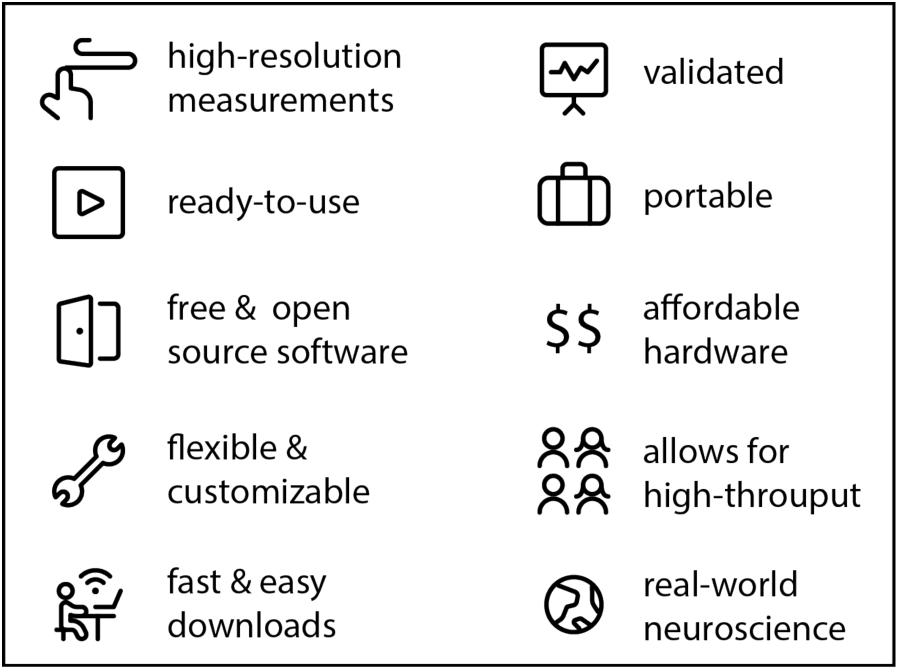
Main advantages of the app for real-world behavioral neuroscience. Our app is ready to use, free, open source, and designed to be flexibly customizable for users who are not programming experts. The app and the data collected can easily be downloaded via USB or email. We include scripts not only for re-programming the app but also for data analyses. Our experimental protocol has been designed for simultaneous high-throughput data collection with several tablets, and validated to reproduce state-of-the-art laboratory results in the real world.

Our custom-made app running on a commercial tablet is a sweet spot between the precision of laboratory equipment and the usability of mobile devices. It is a fact that tinkering with the source code requires some considerable programming knowledge. Yet, our goal here has been to design the code to significantly simplify the task of creating a movement-recording app, specially in comparison with creating it from scratch. Additionally, we have solved some more involved technical issues regarding the access of maximal temporal resolution of touch events and maximal rate of screen refreshing during experiments. In sum, we expect our app (and modifications of it) not only to be usable but actually used.

Broadening the scope, we hope that the methods presented here will be of value to study motor control phenomena outside the laboratory. This may include educational programmes at schools, improving health in hospitals, scientifically studying artistic practices, and even recreational purposes at home. Neuroscience has needed a considerable amount of time to realize the imperative to go “out of the head”. Conceding a certain dose of behavioral “chauvinism” in the face of 21^st^ century “neuralism” (**Gomez-Marin, 2017**), one must take seriously the idea that in order to understand how the brain works we must also ask what it is for (aka, behavior). At the end of the day, everyone willing to spend some time with our App scripts and with 200$ to spend on a tablet can now do high-resolution human behavioral science of laboratory-quality in the real world. The time is ripe to move “out of the lab”.

## Supplementary material

All codes and data used in this study are available. The app in release form (*app-release.apk*) and the source codes to edit it can be obtained from the authors directly in the following online repository: https://github.com/adam-matic/KinematicCognition. The raw behavioral data used in this study together with the scripts used to analyze it in Python within a Jupyter notebook can be found here: https://github.com/adam-matic/KinematicCognition-Analysis.

## Contributions

Idea and conceptualization: AGM; experimental design: AM and AGM; app development: AM; experiments: AM; data analysis: AM; figures: AM and AGM; first manuscript draft: AM, final manuscript: AGM.

## Acknowledgements

We thank María Regina Zaghi Lara, Roberto Morollón, and María del Carmen Lillo Navarro for valuable suggestions on the experimental protocol and for help in testing the app.

## Funding

The authors declare no competing financial interests. The work was supported by the Spanish Ministry of Science (grant BFU-2015-74241-JIN to AGM; pre-doctoral contract BES-2016-077608 to AM) and by the Severo Ochoa Center of Excellence programs (SEV-2013-0317 start-up funds to AGM).

